# Ancient DNA reveals early use of melons in China’s Song Dynasty

**DOI:** 10.64898/2026.02.10.704887

**Authors:** Nathanael Walker-Hale, Yunfei Zheng, Marion Verdenaud, Lara Pereira, Oscar Alejandro Pérez-Escobar, Michaela Preick, Abdelhafid Bendahmane, Michael V. Westbury, Michael Hofreiter, Hanno Schaefer, Xinyi Liu, Guillaume Chomicki, Susanne S. Renner

## Abstract

Melon (*Cucumis melo* L.) domestication is thought to have occurred independently once in Northeast Africa and twice in India, but archaeobotanical seed remains point to a possible additional domestication in China. Because Cucumis seeds are difficult to diagnose morphologically, genomic data from archaeological material are needed to evaluate these scenarios and reconstruct ancient melon traits. We sequenced two Song Dynasty melon seeds from Shuomen Gugang (China), recovering 4.6× and 1.7× nuclear genome coverage. Nuclear and chloroplast analyses place both seeds within cultivated *C. melo* from China, supporting introduction from the broader Asian domestication pool rather than independent Chinese domestication. To assess whether these seeds carried traits associated with sweet dessert melons, we examined loci underlying fruit phenotypes. Neither seed carried alleles for orange flesh; one harbored an allele linked to yellow/orange peel, the other possessed alleles associated with green flesh and reduced acidity. Since wild melons are monoecious, the presence of the derived andromonoecy allele in one seed, associated with rounder fruit shape, suggests early selection on fruit morphology. Together, these findings indicate that Song Dynasty melons were likely consumed as fresh or culinary fruits rather than as sweet dessert melons. Their flesh coloration resonates with Song-period aesthetic sensibilities, exemplified by jade-green celadon ceramics frequently crafted in melon-shaped forms. By anchoring East Asian archaeobotanical remains within modern melon genomic variation, this study provides a temporal framework for melon dispersal into China and shows how ancient genomics can illuminate past crop use.

## Main text

Melon (*Cucumis melo* L.) is an important crop from the Cucurbitaceae family, with worldwide cultivation exceeding 41.5 million tons in 2024 (FAO; http://faostat.fao.org/). Melon domestication has been extensively studied over the past decade, yielding valuable insights into the roles of selection and introgression (1–4).

Molecular evidence suggests that melon domestication occurred independently once in Northeast Africa and twice in India (2, 3). Archaeobotanical evidence suggested a potential independent fourth domestication in the lower Yangtze region of China (5, 6). Domestication is difficult to infer from seeds in Cucurbitaceae because *Cucumis* seed characters overlap across domesticated, feral and wild forms, and preservation can obscure diagnostic features (6, 7). Molecular data are needed to confirm ancient seeds’ taxonomic identity and test among alternative domestication scenarios.

The inconsistency between genetic and archaeological evidence points to three possible scenarios: (a) independent domestication of *Cucumis melo* in ancient China, with descendants persisting among modern Chinese varieties; (b) early Cucumis use in East Asia involving a species other than *C. melo*; and (c) independent Chinese domestication of *C. melo* that was subsequently replaced by varieties introduced from India or Africa, leaving no detectable genetic trace today. Given the continuous archaeological record in the Lower Yangtze since 7000 BP, a combined archaeo-genomic approach is timely.

Numerous *Cucumis* seed remains were recovered from archaeological sites across China. We extracted DNA from 24 seeds: 20 from the Nanshan and Tingshan sites in Shaoxing, dating from the Spring and Autumn period (770-476 BC), but we could not recover sufficient endogenous DNA. Four were recovered from Shuomen Gugang (henceforth Gugang), an ancient port at Wenzhou. During the Song Dynasty (960-1279 CE), Gugang served as a major harbor connecting maritime trading systems across the western Pacific and Indian Oceans. Excavations in 2024 revealed an archaeobotanical assemblage comprising abundant rice as well as peach, grape, olive, jujube and *Cucumis* seeds. Tropical fruits including lychee and kiwifruit were also present, attesting to extensive maritime trade connections. Two *Cucumis* seeds from waterlogged deposits dating to the Southern Song provided sufficient endogenous DNA; we radiocarbon dated another seed from the same context to 930-800 cal. BP (c. 1020-c. 1150 CE).

Here, we generate and analyze ancient DNA from archaeological *Cucumis* remains from China to (i) confirm taxonomic identity of ancient melon seeds, (ii) test whether early Chinese material is consistent with independent domestication versus introduction from India, and (iii) evaluate the fruit traits of ancient Chinese melons.

## Results and discussion

We performed low-coverage sequencing of 32 libraries prepared from 24 *Cucumis* seeds recovered from three sites in China, alongside extraction and library blanks (average coverage from unique hits 0.00017x, **Supplementary Information Online**). Two seeds from Gugang (GG1 and GG4) yielded sufficient endogenous DNA for deep sequencing. After quality filtering and duplicate removal, we recovered 32 and 13 million unique fragments aligning to the *C. melo* DHL92 reference genome (1), yielding 4.6× and 1.7× coverage (138× and 64× chloroplast coverage), respectively. Misincorporation patterns confirmed *bona fide* ancient DNA (**Supplementary Information Online, Dataset S1**). We combined our ancient data with extant accessions from Zhao et al. (3) to infer pseudohaploid sequences and genotype likelihoods. Principal components analysis and phylogenetic analysis of four-fold degenerate sites place our ancient seeds closest to cultivated *C. melo* from China in clade III (*agrestis* clade *sensu* ref. 3) (**Fig. 1**), a result consistent with chloroplast data and robust to missing data (**Supplementary Information Online, Dataset S1**). This placement supports introduction from the broader Asian domestication pool rather than independent Chinese domestication.

**Figure 1.**
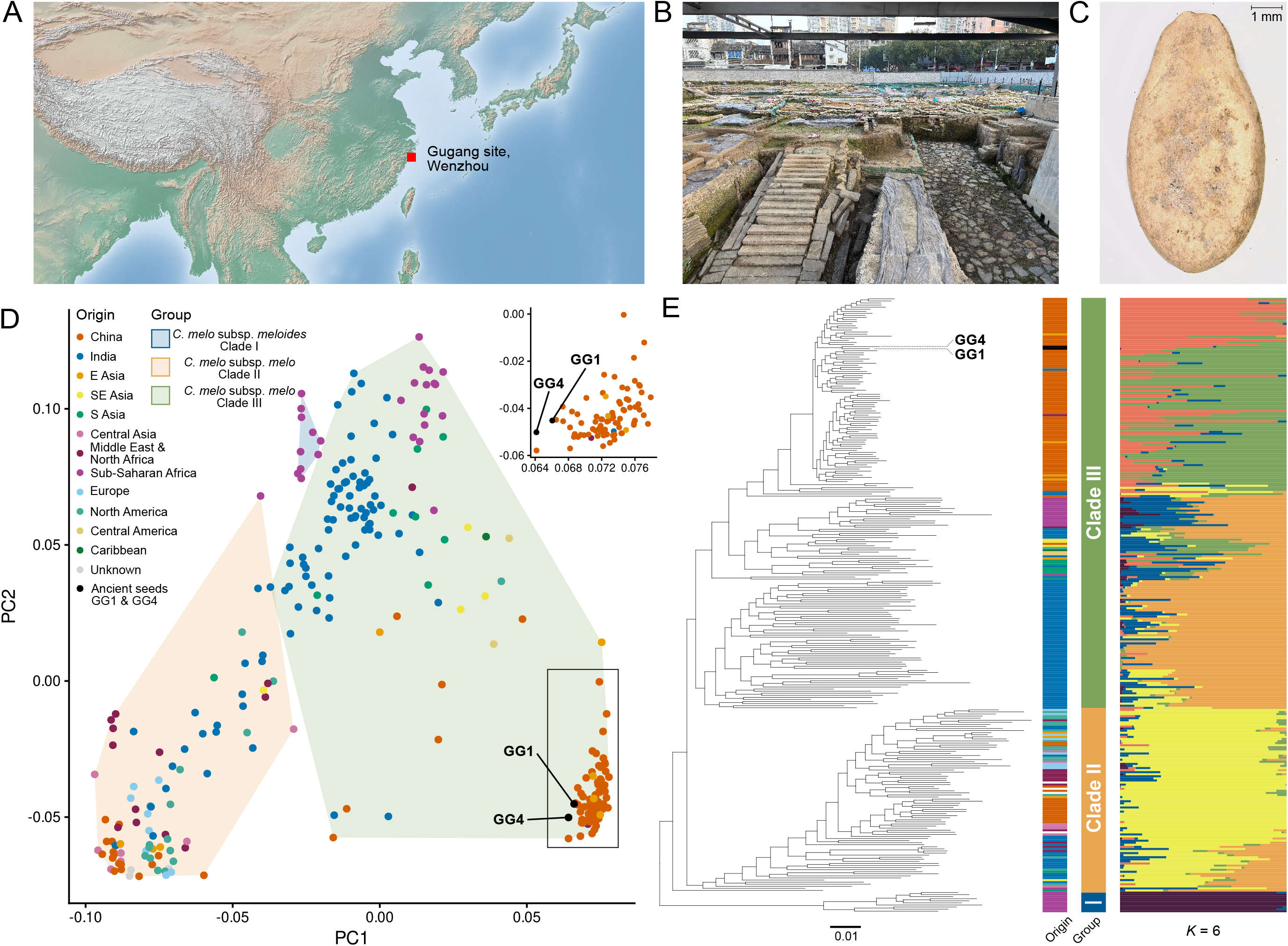
Phylogenomic placement of ancient seeds. (**A**) Location of the Gugang archaeological site in Wenzhou, China. (**B**) The Gugang site. (**C**) Ancient melon seed from Gugang. (**D**) Principal Components Analysis of ancient seed genomes alongside 294 extant melon genotypes, based on inferred genotype likelihoods. (**E**) Maximum likelihood phylogeny of the same genotypes plus 9 outgroups, based on 59,299 four-fold degenerate sites; branch lengths in expected substitutions per site. Tip colors indicate geographic origin and group (inset). Admixture proportions for K = 6 ancestral populations, estimated from genotype likelihoods at positions with no missing data, are aligned to tree tips. For K = 2 to K = 16, see **Supplementary Materials Online**.

The genetically informed identification is consistent with the morphological observation of the waterlogged remains (**Fig. 1C**) and raises questions about the dispersal routes of cultivated *C. melo* originating in Africa and South Asia. Beyond the lower Yangtze, morphologically attested *C. melo* has been reported from ancient burials in northwestern China (c. 700 CE) and the tomb of a Han prince (59 BCE) in southern China, indicating that cultivated *C. melo* was introduced before the Han Dynasty. It is plausible that *C. melo* reached southern China during the Han period via Himalayan routes rather than the northern Silk Route, with subsequent introductions through maritime networks during the Song. The relationship between our Song-dynasty seeds and Neolithic *Cucumis* (c. 7000-4000 BP) in the lower Yangtze remains unresolved; given the continuity of *Cucumis* in archaeobotanical records spanning the Neolithic and Bronze Age, a local domestication scenario from a gene pool not represented by extant genotypes cannot be ruled out. Further aDNA sampling targeting Neolithic or Bronze Age seed remains would be needed to resolve this question.

To infer the potential phenotypes and likely uses of these ancient melons, we compared read alignments at key loci governing fruit traits in *C. melo. CmOR* is involved in β-carotene biosynthesis; the His108 allele conferring orange flesh is dominant over Arg108 conferring white/green flesh (8). Both GG1 and GG4 showed evidence of the Arg108 allele, suggesting they lacked orange flesh; in GG1 the probability of missing a heterozygote given the coverage is very low (**Fig. 2G, H**). *CmRPGE1* (9) encodes a protein involved in plastid development and chlorophyll maintenance; variation at this locus determines whether non-orange melons develop white flesh (chlorophyll loss) or green flesh (chlorophyll retained). We found GG4 harboring the green flesh allele (**Fig. 2H**), consistent with a higher frequency of green flesh in clade III, but due to low coverage we cannot rule out heterozygosity. For peel coloration, we found no truncations in *CmAPRR2*, suggesting dark immature skin (10, **Fig. 2G, H**). GG1 lacked a white-peel-associated deletion in *CmKFB* (11; **Fig. 2G**), suggesting yellow peel. We assessed genotypes at four loci involved with climacteric ripening (12); GG1 and GG4 largely featured genotypes associated with non-climacteric varieties, suggesting weak or no climacteric ripening and consequently green-yellow mature peel in GG1 (**Fig. 2G**). The fruits were likely non-sutured (**Fig. 2G, H**, 3). A single read in GG4 supports a 12-bp duplication in *CmPH* controlling fruit acidity (13), suggesting non-acidic flesh. Additionally, both samples fall in a clade with a high frequency of non-acidic accessions (**Fig. 2D**). We found no loss-of-function truncations in the bitterness regulator *CmBt*, although bitterness may also be regulated by *CmBt* promoter variation (3). Andromonoecy has been favored under domestication because it is associated with rounder fruits and facilitates breeding (14). CmACS7 underlies the transition from monoecious to andromonoecious accessions; a single substitution (Ala57Val) confers this shift (14). GG1 carries the derived Val allele, suggesting it was andromonoecious. Read alignments supporting genotypes for all traits are shown in **Dataset S1**.

**Figure 2.**
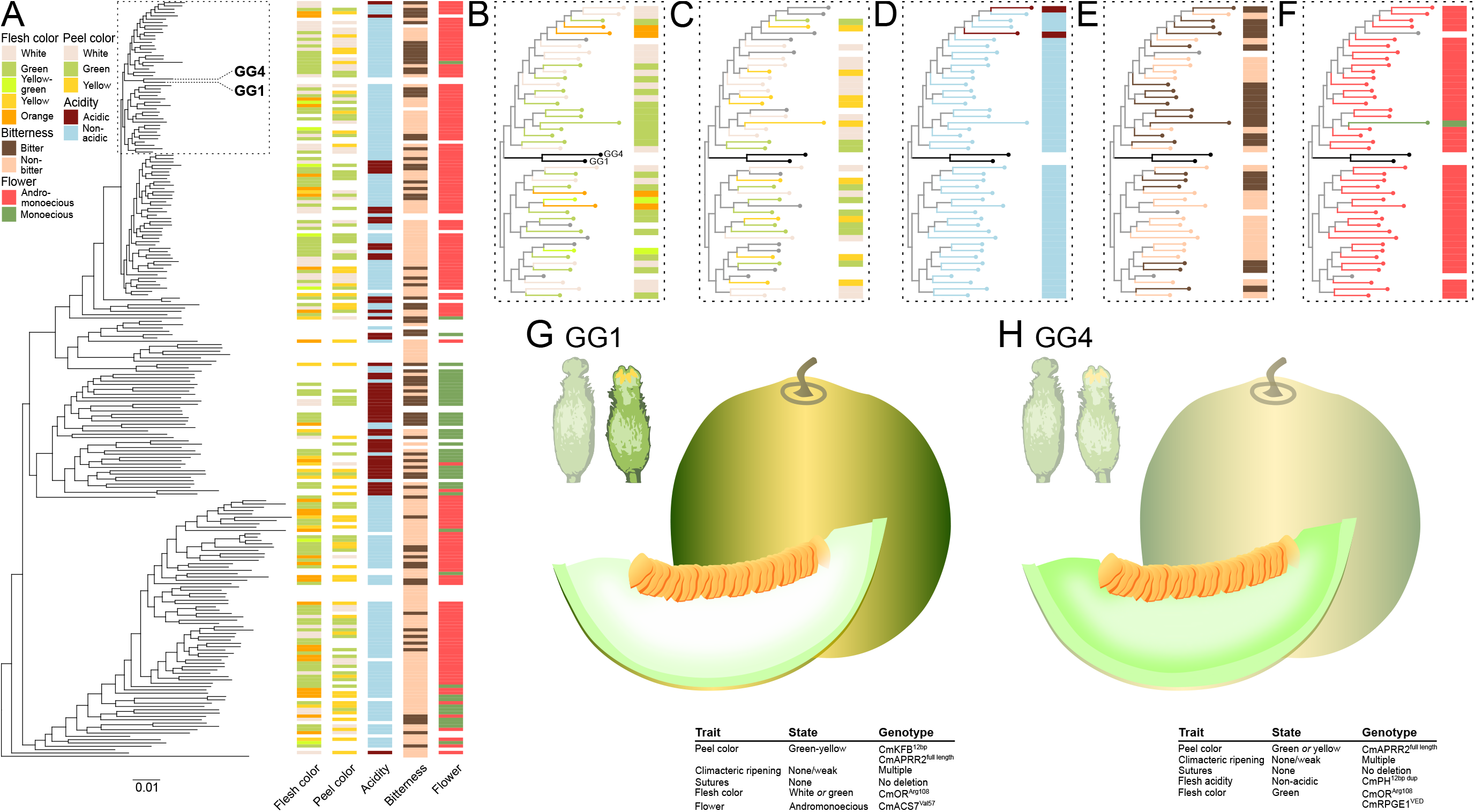
Comparative analysis of ancient fruit phenotypes. (**A)** Distribution of four phenotypes across tips in the inferred tree; samples lacking all trait data were removed. (**B-E)** Subtrees surrounding our ancient samples for (**B**) fruit flesh color, (**C**) fruit peel color, (**D**) fruit acidity, (**E**) fruit bitterness, (**F**) flower type. (**G-H**) Reconstructed appearance of ancient fruits and flowers based on inferred phenotypes. Tables summarize trait inferences and supporting genotypes; traits not inferred are shown with reduced opacity. Although exact fruit shape and size cannot be inferred, the CmACS7^Val57^ allele is associated with rounder fruits. Not drawn to scale.

Several extant melons combine the same trait suite inferred from our Song Dynasty seeds: non-orange flesh, often yellow peel, and sometimes white to light-green flesh. These are commonly used as fresh, mild fruits, vegetables or seeds. Analogous varieties include Korean oriental melon (*makuwa*), Japanese pickling melon (*conomon*), South Indian culinary melon (*acidulus*), and Sudanese tibish and seinat (4). Taken together, these modern analogues suggest that melons sharing the trait combination observed in our ancient samples (non-orange, white or potentially green flesh, and green-yellow peel) were likely used in Song Dynasty China as crisp, mildly sweet fruits, in fresh or processed forms, and potentially also as a source of edible seeds, rather than being selected exclusively for high sugar content. This interpretation is further supported by cultural considerations: “oriental melon” types with light-green flesh and yellow peel have long held symbolic significance in East Asia, where abundant seeds were associated with fertility and continuity, and green flesh resonated with established aesthetic values. In particular, green celadon ceramics produced in the Lower Yangtze, including Longquan celadon, were frequently fashioned in melon-shaped forms, with glazes echoing the jade-green tones of melon flesh, a typology that became especially prominent during the Song period.

## Materials and Methods

We selected 24 *Cucumis* seed remains from three archaeological sites in China for ancient DNA analysis. DNA was extracted following established protocols for degraded archaeobotanical material, and single-stranded Illumina sequencing libraries were prepared in a dedicated ancient DNA facility with negative controls at all steps. Two seeds from Shuomen Gugang (GG1 and GG4) yielded sufficient endogenous DNA; deep sequencing recovered 4.6× and 1.7× nuclear genome coverage, respectively. We combined these ancient data with 305 extant *C. melo* accessions and performed pseudohaploid genotype inference following (15), phylogenomic analysis of four-fold degenerate sites, principal components analysis, and admixture analysis to place the ancient seeds within the context of modern melon diversity. We further inspected read alignments at loci governing fruit flesh color, peel color, acidity, bitterness, suture, climacteric ripening, and sex determination to infer ancient fruit phenotypes. Full methods, including details of read processing, error assessment, and all analytical parameters, are provided in the ***Supplementary Information Online***.

## Acknowledgments

We thank the UK Natural Environment Research Council for an Independent Research Fellowship (NE/S014470/3) and ERC/UKRI for a frontier research grant (EP/X026868/1) to G.C. A.B. is funded by NectarGland ERC Project (101095736). We thank Melissa Ritchey for assistance with imaging the archaeological seed remains. We thank Jordi Garcia-Mas for reading an earlier version of this manuscript.

## Data availability

Datasets S1 is available here in Figshare: https://doi.org/10.6084/m9.figshare.31830979. Raw reads have been deposited on the NCBI under PRJNA1422990.

## Author Contributions

S.S.R., G.C., N.W.-H., and X.L. designed research; N.W.-H. performed research, with support from M.P., M.V.W, L.P., O.A.P.-E.; Y.Z. and X.L. provided the ancient seeds, M.P. and M.H. provided ancient DNA extraction facilities; M.V.W. provided sequencing facilities; A.B., H.S., and X.L. contributed analytical tools and expertise; N.W.-H., G.C., and S.S.R. wrote the paper with input from all authors; G.C. secured funding.

