## Supporting Information for "Ancient DNA reveals early use of melons in China’s Song Dynasty"

**This PDF file includes:**

Supplementary Materials and Methods

SI References

**Supplementary Materials and Methods**

### *Ancient DNA extraction, sequencing, and radiocarbon dating*

We selected 24 *Cucumis* seed remains to attempt ancient DNA analysis: 20 from the Nanshan (NS) and Tingshan (TS) sites in Shaoxing, dating from the Spring and Autumn period (770–476 BC), and four from Shuomen Gugang (GG), an ancient port at Wenzhou dating from the Song Dynasty (960–1279 CE). Seeds were stored at 4°C prior to DNA extraction. The outer seed coat (testa) was separated from the inner tissues and used separately for extraction.

DNA was extracted from the seed coats and inner parts of GG1–4 and NS1–4, and seed coats of NS 5–10 and TS 1–10 following Dabney et al. (1) and Wales et al. (2). For digestion, a lysis buffer containing 0.5% (w/v) *N*-lauroylsarcosine (Sigma-Aldrich L9150-50G), 50 mM Tris-HCl (Thermo Fisher Scientific 15568025), 20 mM EDTA (VWR E177-500MLDB), 150 mM NaCl (Thermo Fisher Scientific AM9760G), 3.3% 2-mercaptoethanol (Sigma-Aldrich 63689-25ML-F), 50 mM DL-dithiothreitol (Sigma-Aldrich D9779-250MG), and 0.25 mg/mL proteinase K (Promega V3021) was applied to the powdered material as described in Wales et al. (2). DNA purification followed Dabney et al. (1) with reduced centrifugation speed (450 × g).

Library construction and sequencing were performed in the ancient DNA facility at the University of Potsdam; negative controls were included at all steps. DNA extracts were converted to Illumina sequencing libraries using the single-strand approach of Gansauge et al. (3) using custom adapters from Gansauge and Meyer (4). Quantitative PCR (qPCR) was performed on a PikoReal 96 Real-Time PCR machine (Thermo Fisher Scientific TCR0096) using 0.2% of each unamplified library with the following thermal profile: 10 min initial denaturation at 95°C, followed by 40 cycles of 15 s at 95°C, 30 s at 60°C, and 1 min at 72°C. Each 10 µL qPCR reaction contained 1 µL diluted library, 1× SYBR Green qPCR Master Mix (Applied Biosystems 4309155), and 0.5 µM each of primers IS7 and IS8; three technical replicates were run per library. Indexing PCR was performed using the cycle number determined by qPCR, adding 8-bp indices to both 5′ and 3′ adapters. Preliminary DNA sequencing was performed on an Illumina NextSeq 500 sequencing platform, using the 500/550 High Output v2 kit (75 cycles, Illumina FC-404-2005), with a custom read-1 (4) and a custom index-2 (5) sequencing primer at the University of Potsdam, Germany. For deep sequencing of two successful seeds, pooled libraries (5:4 ratio) were sequenced on an Illumina NovaSeq S2 flowcell (two lanes, 100 cycles), using the same sequencing primers as above, at the Globe Institute, University of Copenhagen, Denmark.

All but two of these seeds (GG1 and GG4) did not yield sufficient endogenous DNA for deep sequencing. Both successful seeds derive from contexts dated to the Southern Song period (1127–1279 CE). We used another *Cucumis* remain (GG6) from the same context for radiocarbon analysis at Center for Applied Isotope Studies, University of Georgia, USA. The sample preparation followed well-established acid-base-acid (ABA) chemical pre-treatment, followed by combustion and graphitization prior to accelerator mass spectrometry (AMS) measurement. The result was calibrated by the OxCal (v4.4) and the IntCal20 calibration curve. The result is 928–796 cal. BP (reported relative to 1950 CE at the 95.4% probability range), corresponding to approximately 1020–1150 CE, placing it securely within the Southern Song period. GG1 and GG4 were not directly dated but derive from the same stratigraphic unit.

### *Ancient DNA read processing*

Raw reads from the two ancient seeds were preprocessed and aligned using the BAM pipeline in PALEOMIX v1.3.10 (6). AdapterRemoval v2.3.4 (7) was used to remove adapter sequences, with a maximum mismatch threshold of 1/3, trimming Ns and low-quality bases and discarding fragments shorter than 30 bp. Trimmed reads were aligned to the *C. melo* DHL92 v4.0 nuclear reference (8) and a chloroplast reference (NC_015983.1; 9), from which one copy of the inverted repeat was removed, using BWA v0.7.18 (10) with the backtrack algorithm, which is appropriate for short reads. To assess characteristic patterns of post-mortem DNA damage, we estimated the proportion and position of misincorporations in aligned reads using mapDamage v2.2.2 (11). Based on these results, we trimmed the first and last three nucleotides from each read and removed fragments shorter than 30 bp, prior to realignment with BWA backtrack. Quality scores were then rescaled to account for post-mortem damage using mapDamage, reads were realigned around indels with GATK v3.8.1 (12), and duplicates were marked with Picard MarkDuplicates implemented in GATK v4.6.2.

### *Raw read data processing*

Sequencing data for 1,175 accessions of *Cucumis melo* (subsp. *melo* and *agrestis*) from Zhao et al. (13) were downloaded from NCBI (BioProject PRJNA565104), and inbred lines were excluded. Cultivated accessions were further subsampled, yielding a final dataset of 305 samples. Read quality was assessed with FastQC v0.12.1 (14), and Ns and low-quality bases were removed with Trim Galore! v0.6.10 (https://www.bioinformatics.babraham.ac.uk/projects/trim_galore/) using Cutadapt v5.2 (15). Reads were aligned to the same references as above using BWA-MEM (16) and similarly subjected to indel realignment and duplicate marking.

### *Ancient DNA pseudohaploidization and error assessment*

Due to low coverage, we employed a pseudohaploid approach following Pérez-Escobar et al. (17): sample depths and nucleotide occurrences were calculated using ANGSD v0.940 (-doCounts 1 -minQ 30 -minMapQ 20 -dumpCounts 3). Pseudohaploid sequences for the chloroplast genome and each of the 12 nuclear chromosomes were inferred using Consensify v2.4.0, with maximum depth set to the upper 95th percentile of coverage for each sample (18).
We assessed error rates by computing the excess of derived alleles in ancient relative to extant samples using the -doAncError function in ANGSD, with a modern *C. melo* subsp. *agrestis* consensus sequence as reference (17). We created a reference consensus FASTA from the MS-996 BAM with ANGSD (-doFasta 2 -doCounts 1 -minQ 30 -minMapQ 20 -setMinDepth 15), then used this together with the reference DHL92 genome to calculate excess substitutions as error rates for each ancient sample and three high-coverage accessions (MS-1001, MS-1002, and MS-1021) as controls, using ANGSD (-doAncError 1). We further compared the transition/transversion ratio (Ts/Tv) against the

reference for positions genotyped in our ancient pseudohaploid sequences and positions in ANGSD consensus FASTAs (-doFasta 2 -minQ 30 -minMapQ 20 -setMinDepth 5) from 17 high-coverage accessions from the Zhao et al. (13) dataset. We generated BED files of the non-gap positions in each ancient pseudohaploid FASTA and extracted the same positions from each extant FASTA with bedtools getfasta in bedtools v2.31.1 (19).

### *Phylogenomic placement of ancient samples*

We used the pseudohaploid sequences from our ancient samples for phylogenomic inference. For extant accessions, consensus sequences were generated at each position meeting quality and depth thresholds (-doFasta 2 -minQ 30 -minMapQ 20 -setMinDepth 5).
The chloroplast alignment was analyzed in a maximum likelihood framework using IQ-TREE v3.0.1 under a GTR+F+G model with 1,000 ultrafast bootstrap replicates and 1,000 replicates for the SH-aLRT test (-m GTR+F+G -B 1000 -alrt 1000; 20, 21). For nuclear data, four-fold degenerate sites were identified using degenotate v1.2.4 (22) and extracted from each pseudohaploid genome. Invariable columns were removed with goalign v0.4.0 (--char MAJ --cutoff 1; 23), and columns with >25% gaps were filtered using pxclsq from phyx (24). The final nuclear alignment was analyzed with IQ-TREE using the same settings as above, except that we added an ascertainment bias correction to account for the removal of fixed sites (-m GTR+F+G+ASC).

### *Population structure of* Cucumis melo

To call genotype likelihoods, we used ANGSD with the GATK model, applying the default minimum Minor Allele Frequency cutoff of 0.05 and testing for polymorphism with a SNP p-value cutoff of $1\times10^{-6}$, using the same quality thresholds as previously (-GL 2 -doMajorMinor 1 -doMaf 1 -doGlf 2 -SNP_pval 1e-6 -minQ 30 -minMapQ 20 -minMaf 0.05). To account for missing data, we repeated this process requiring that all individuals be present (-minInd 296). For Principal Components Analysis (PCA), we used pcangsd v1.36.4 (25) to calculate the covariance matrix, setting a maximum of 1,000 iterations (--iter 1000). PCA on the resulting matrix was conducted in R v4.5.0 (26) using the eigen() function. For both the full and reduced datasets, the number of eigenvectors automatically selected by pcangsd was 15, but the full matrix was unable to converge. We therefore present results from the reduced dataset here. In order to further account for the potential influence of missing data on our inferences, we conducted PCA with EMU v1.6.0 (27). To do this, we generated pseudohaploid genotype calls with ANGSD (-doHaploCall 2 -doIBS 2 -doCov 1 -doGlf 2 -doMajorMinor 1 -GL 2 -doCounts 1 -doMaf 1 -SNP_pval 1e-6 -minMaf 0.05) and converted them with plink v1.90 (28). EMU was run with 15 eigenvectors (-e 15). To calculate admixture proportions, we used NGSadmix v33 (29), setting the likelihood difference in 50 iterations to 0.05. For consistency with pcangsd, we analyzed K=2–16, and used evalAdmix v0.962 (30) to assess the residual correlations for each K. We assessed a scree plot of the sum of absolute values of correlations and picked K=6. Different admixture components were aligned across runs with CLUMPAK (31).

### *Comparative analysis of functional variants*

Using IGV (32), we manually inspected the alignments of our ancient DNA reads at loci known to determine fruit phenotypes in *Cucumis melo*, namely the bHLH regulator of bitterness CmBt (33), the fruit color determinant CmOr (34), acidity regulator CmPH (35), peel color regulator CmAPRR2 (36) and CmKFB (37), the ~1 Kb deletion associated with sutured fruits (13), and the recently described locus determining white or green flesh in non-orange varieties, CmRPGE1 (38). We also assessed the regulator of andromonoecious flowering, CmACS7, for the presence of Ala57Val controlling transitions from monoecy to andromonoecy (39). For this analysis, we attempted to increase informative data by reducing our fragment length filter to a minimum of 25 bp after trimming each end.

We also assessed variation at four loci associated with climacteric ripening behavior: CmCTR1 and CmROS1 (40), *CmNAC-NOR* (41), and *CmERF024* (42). For this analysis, because key variants are not fully described and due to the preponderance of small indels at these loci, we first called genotypes with GATK v4.6.2 (HaplotypeCaller --min-pruning 1 --min-dangling-branch-length 1 --active-probability-threshold 0.001, genotyping with GenotypeGVCF after importing with GenomicsDBImport). As controls for this analysis, we used a sample of 20 accessions each from clade II and III likely to show climacteric ripening (*cantalupensis*, *momordica*) or non-climacteric ripening (*inodorus*, *conomon*). We removed any positions not covered by at least two reads in either ancient sample, and removed positions with MAF < 0.05. Finally, we conducted a 2×N chi-squared test on the allele frequencies at each position against climacteric or non-climacteric behavior, using a chi-squared test in R with p-values from 10,000 simulations, and assessed the overlap between our ancient seed genotypes and genotypes enriched in each ripening behavior at significant sites. Although this probably overstates the evidence for association due to many genotypes in the same gene likely coming from shared haplotypes, it provides a coarse and effective way to assess putative ripening behavior in our ancient accessions.
