## Supplementary material for "Ancient DNA reveals early use of melons in China’s Song Dynasty": Datasets S1

**Table S1: Sequencing results.** Reads gives the number of output reads from sequencing, retained reads' gives the number after removing adapters, Ns, quality trimming, and removing reads shorter than 30bp. 'Mapped reads' is the number of retained reads mapping to the *C. melo* nuclear reference. 'Unique reads' is the number of remaining reads after marking and filtering duplicates. 'Unique fraction' is the proportion of retained reads that are unique reads. 'Coverage' is the average coverage of the nuclear genome from unique reads. Note that GG1-shell_final and GG4-shell_final are the two deep sequencing runs, and their filtering includes removing min 30bp reads after trimming the first and last 3bp from each read.

| sample | reads | retained reads | mapped reads | unique reads | unique fraction | coverage |
|---|---|---|---|---|---|---|
| GG1-shell | 458391 | 300511 | 25753 | 25246 | 0.08401 | 0.0036082229 |
| GG1-inner_part | 362359 | 15962 | 231 | 229 | 0.01435 | 2.41E-05 |
| GG2-shell | 384011 | 170925 | 453 | 442 | 0.00259 | 5.46E-05 |
| GG2-inner_part | 331517 | 41014 | 34 | 34 | 0.00083 | 3.68E-06 |
| GG3-shell | 348168 | 58565 | 151 | 147 | 0.00251 | 1.51E-05 |
| GG3-inner_part | 291719 | 31487 | 21 | 21 | 0.00067 | 2.18E-06 |
| GG4-shell | 554064 | 398696 | 7484 | 7358 | 0.01846 | 0.0009952991 |
| GG4-inner_part | 210653 | 14884 | 71 | 71 | 0.00477 | 7.19E-06 |
| NS1-shell | 606752 | 356955 | 44 | 42 | 0.00012 | 4.72E-06 |
| NS1-inner_part | 557727 | 68743 | 23 | 20 | 0.00029 | 2.14E-06 |
| NS2-shell | 534321 | 201840 | 63 | 62 | 0.00031 | 7.28E-06 |
| NS2-inner_part | 450671 | 116501 | 28 | 26 | 0.00022 | 2.72E-06 |
| NS3-shell | 507685 | 266588 | 28 | 27 | 0.00010 | 3.03E-06 |
| NS3-inner_part | 429269 | 90577 | 34 | 31 | 0.00034 | 3.14E-06 |
| NS4-shell | 607044 | 74543 | 161 | 161 | 0.00060 | 2.01E-05 |
| NS4-inner_part | 434989 | 269750 | 30 | 29 | 0.00039 | 2.92E-06 |
| NS5 | 930440 | 354038 | 223 | 219 | 0.00062 | 2.45E-05 |
| NS6 | 819992 | 371377 | 72 | 70 | 0.00019 | 6.86E-06 |
| NS7 | 601152 | 216641 | 115 | 101 | 0.00047 | 1.03E-05 |
| NS8 | 641125 | 237256 | 85 | 78 | 0.00033 | 8.42E-06 |
| NS9 | 926206 | 218242 | 138 | 123 | 0.00056 | 1.23E-05 |
| NS10 | 998743 | 513556 | 126 | 117 | 0.00023 | 1.27E-05 |
| TS1 | 727029 | 575763 | 371 | 365 | 0.00063 | 4.72E-05 |
| TS2 | 941773 | 586294 | 813 | 801 | 0.00137 | 9.95E-05 |
| TS3 | 868115 | 633789 | 587 | 569 | 0.00090 | 7.13E-05 |
| TS4 | 948912 | 500797 | 970 | 950 | 0.00190 | 0.0001155136 |
| TS5 | 754545 | 544223 | 472 | 459 | 0.00084 | 6.62E-05 |
| TS6 | 824101 | 619553 | 320 | 313 | 0.00051 | 4.14E-05 |
| TS7 | 601686 | 370255 | 384 | 378 | 0.00102 | 4.71E-05 |
| TS8 | 528899 | 327966 | 426 | 425 | 0.00130 | 5.61E-05 |
| TS9 | 782212 | 624486 | 337 | 331 | 0.00053 | 4.54E-05 |
| TS10 | 733963 | 568288 | 501 | 493 | 0.00087 | 6.30E-05 |
| GG1-shell_final | 2276323175 | 1195716243 | 87138654 | 32609746 | 0.02727 | 4.5642467852 |
| GG4-shell_final | 2595936529 | 1522189954 | 26204506 | 13475749 | 0.00856 | 1.7145175764 |

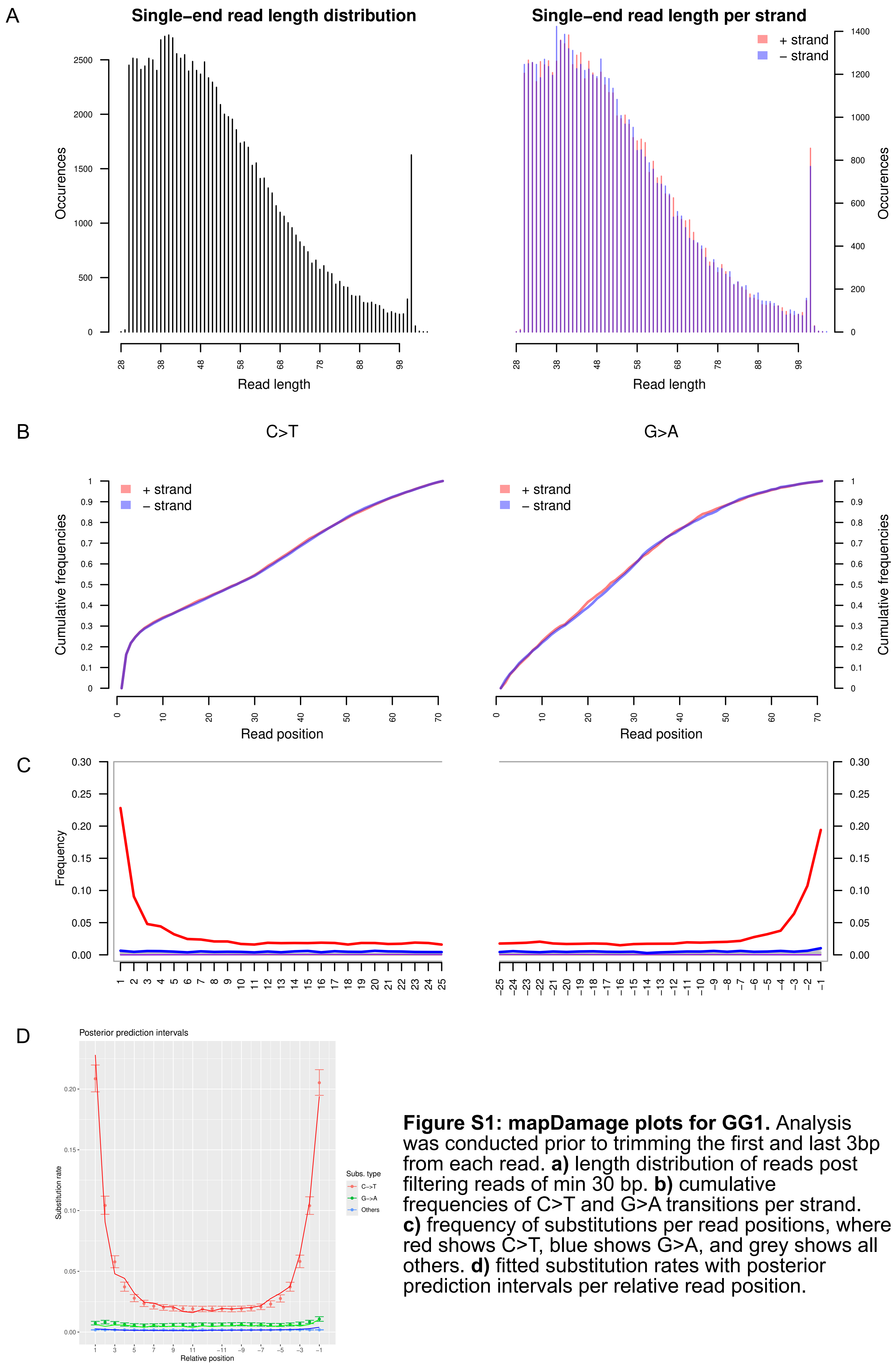


**Figure S1: mapDamage plots for GG1.** Analysis was conducted prior to trimming the first and last 3bp from each read. **a)** length distribution of reads post filtering reads of min 30 bp. **b)** cumulative frequencies of C>T and G>A transitions per strand. **c)** frequency of substitutions per read positions, where red shows C>T, blue shows G>A, and grey shows all others. **d)** fitted substitution rates with posterior prediction intervals per relative read position.

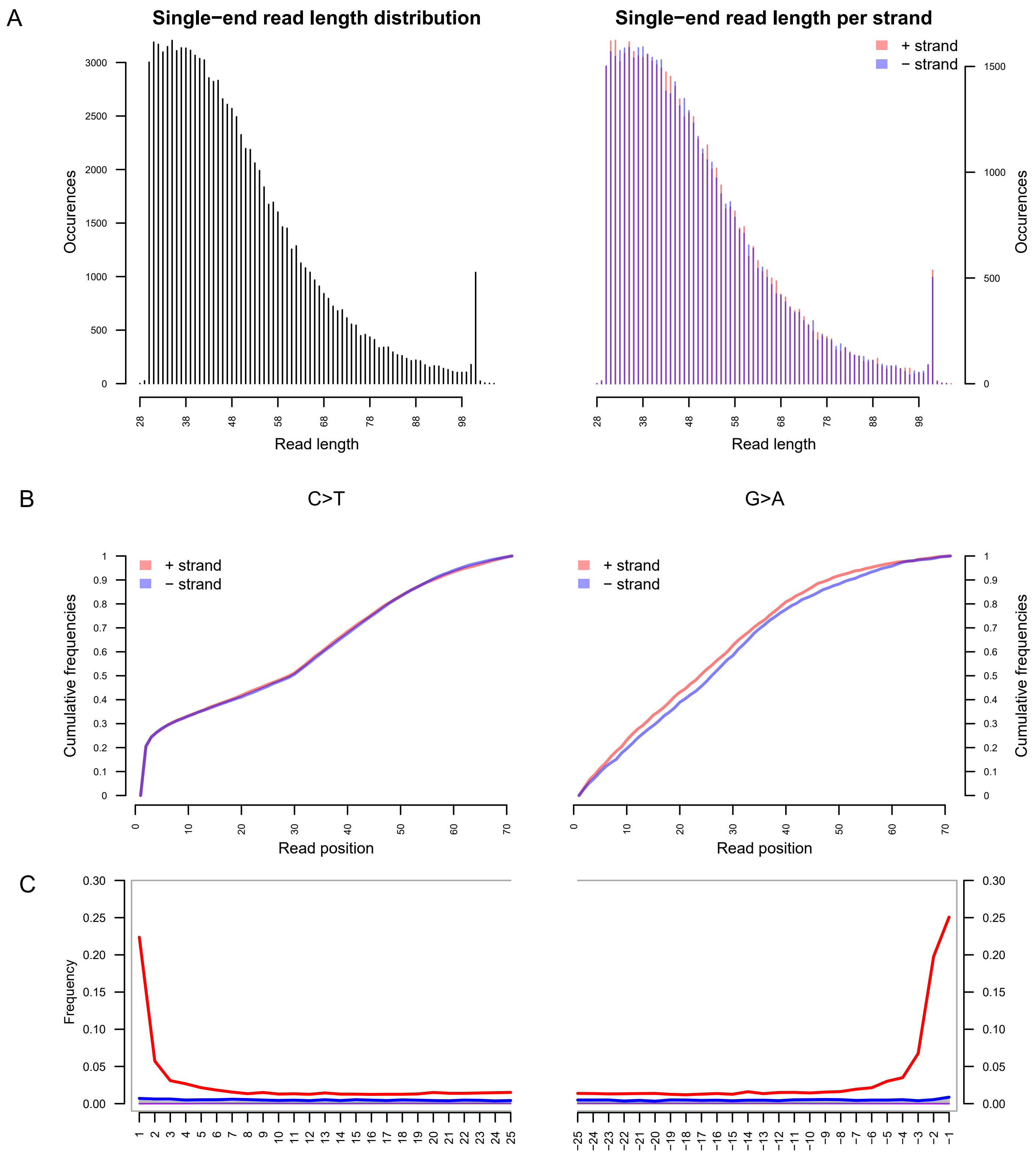


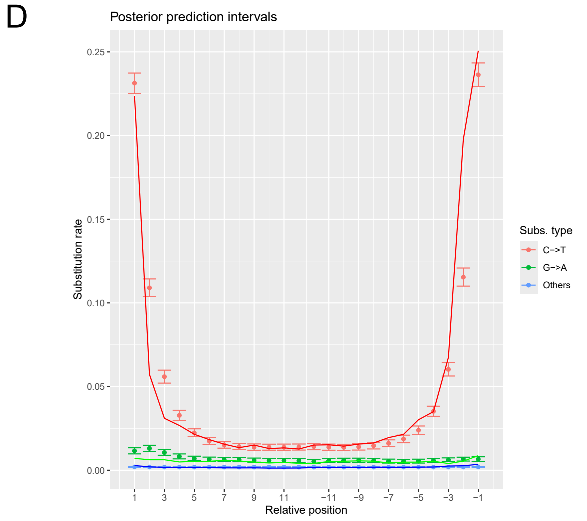


**Figure S2: mapDamage plots for GG4.** Analysis was conducted prior to trimming the first and last 3bp from each read. **a)** length distribution of reads post filtering reads of min 30 bp. **b)** cumulative frequencies of C>T and G>A transitions per strand. **c)** frequency of substitutions per read positions, where red shows C>T, blue shows G>A, and grey shows all others. **d)** fitted substitution rates with posterior predictionintervals per relative read position.

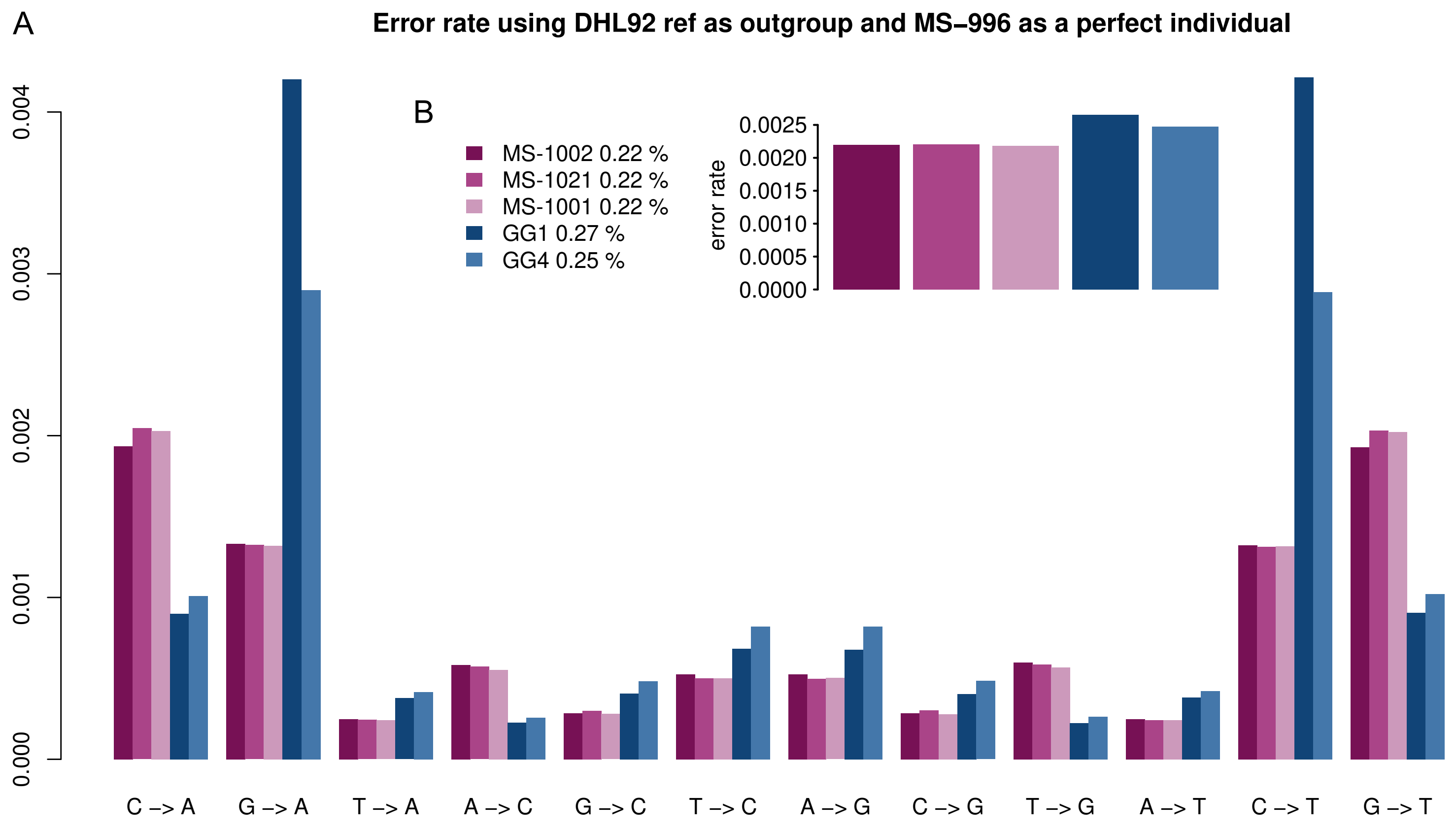


**Figure S3: Error rates of ancient genomes estimated using ANGSD's 'perfect genome' approach.** The DHL92 reference was used as an outgroup and the 22.5x accession MS-996 was used as the perfect individual. Ancient genomes were compared to three other individuals with relatively high coverage as controls. **a)** error rate estimated over multiple substitution types. **b, inset)** overall error rate. Note that this analysis does not take into account the mapped strand, explaining the presence of G>A transitions as well as C>T transitions, contrary to the findings in **Figs. S1** and **S2**.

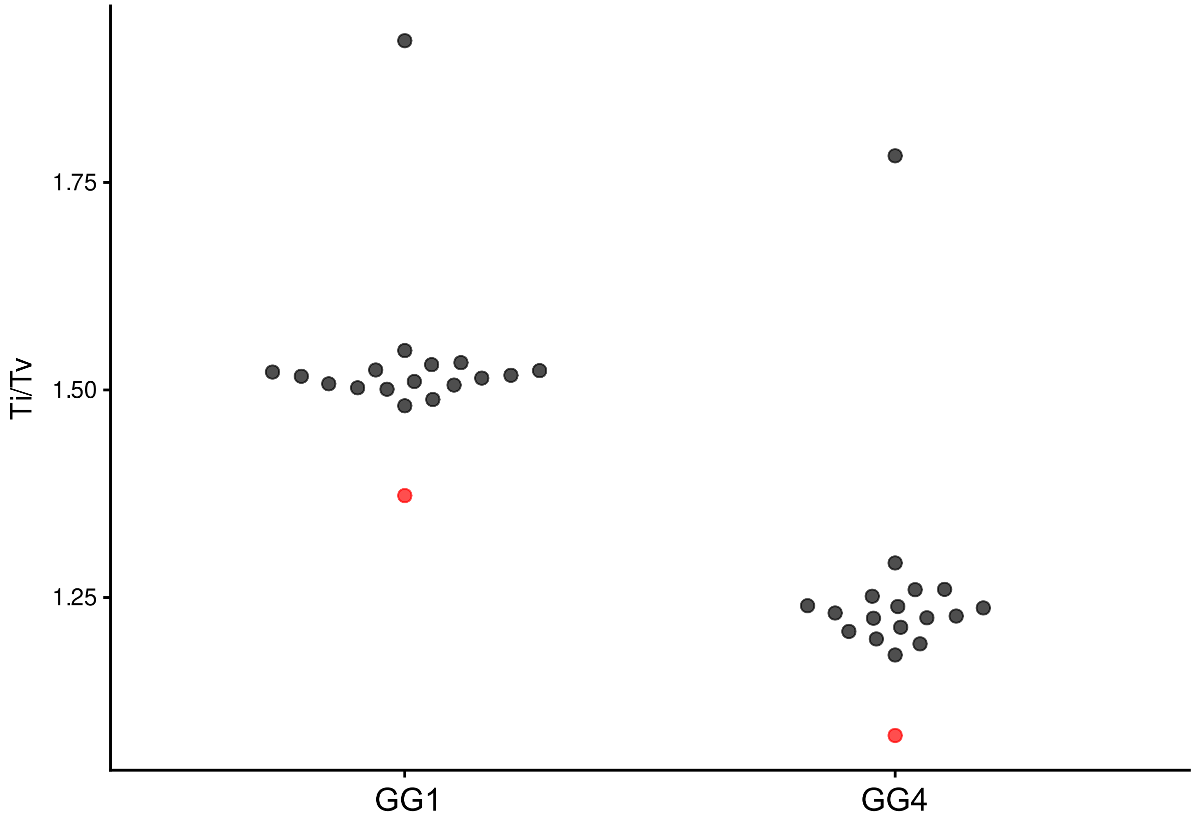


**Figure S4: Transition/Transversion ratios between extant samples and ancient samples.** Transition/Transversion ratios measured between each pseudohaploid ancient genome or each consensus extant genome, and the reference. For each of the ancient seeds, only positions that were present in each pseudohaploid genome were considered. Red points show the ancient seed ratio. The extant samples used for this comparison are: MS-1000, MS-1001, MS-1002, MS-1006, MS-1016, MS-1020, MS-1021, MS-1025, MS-983, MS-988, MS-990, MS-991, MS-993, MS-996, MS-997, MS-998 and MS-999.

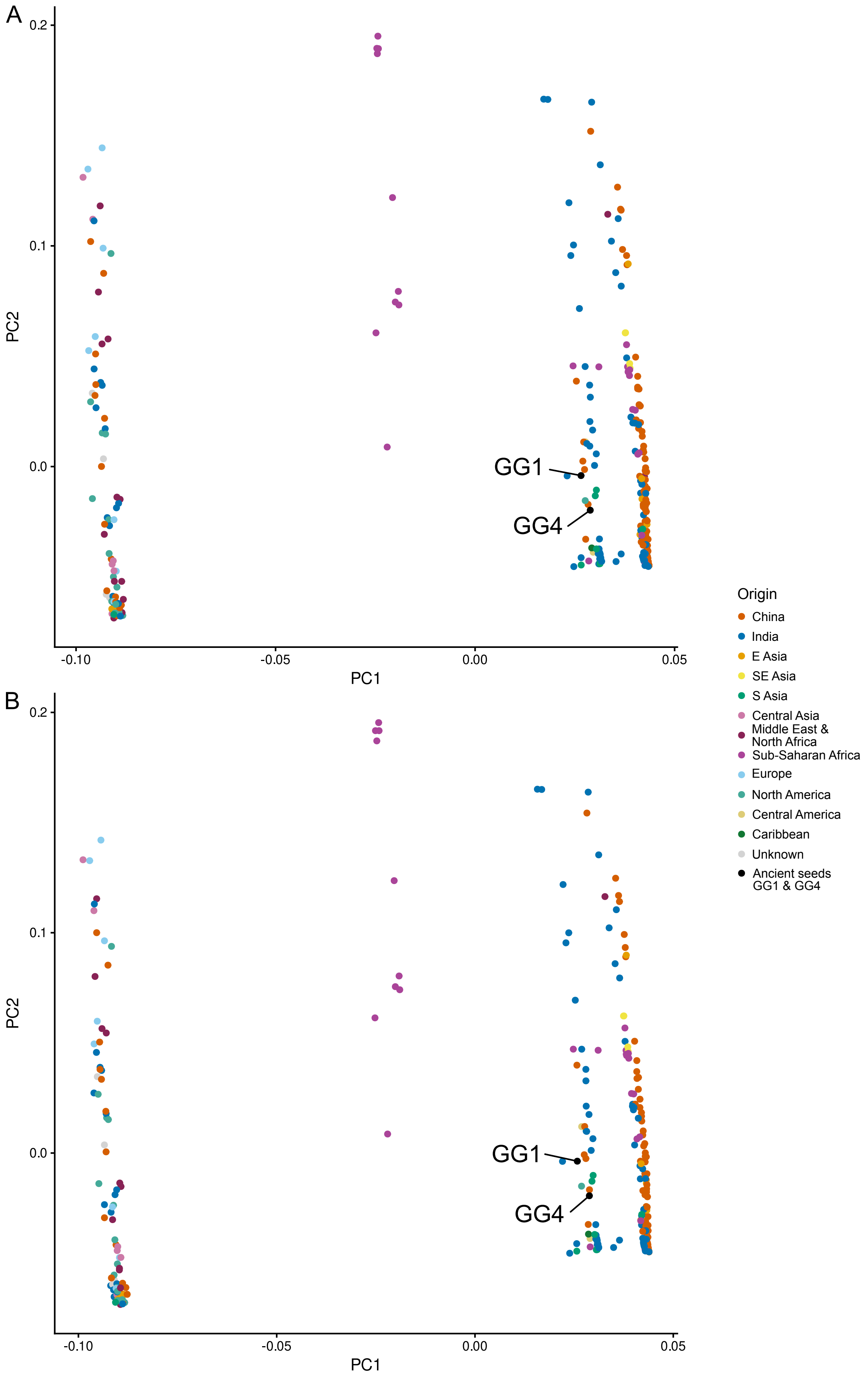


**Figure S5: PCA of chloroplast genotype likelihoods** from **a)** all sites **b)** sites present in all 296 individuals. Points are coloured by geographic origin of the accession and the two ancient seed genomes are labelled.

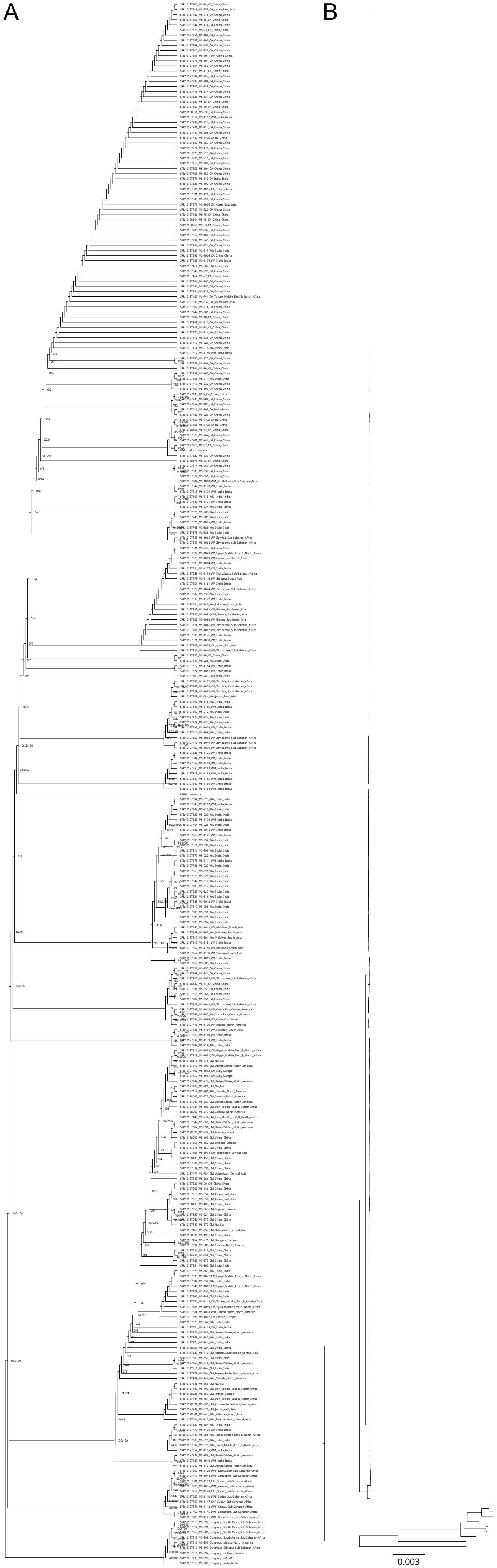


**Figure S6: Chloroplast phylogenetic tree. a)** cladogram. Node labels give support values from SH-aLRT/UFBoot. **b)** phylogram with branch lengths in expected substitutions per site. Note that many  branches are length 0 due to identical sequences in consensus. Scale bar gives 0.003 expected substitutions per site.

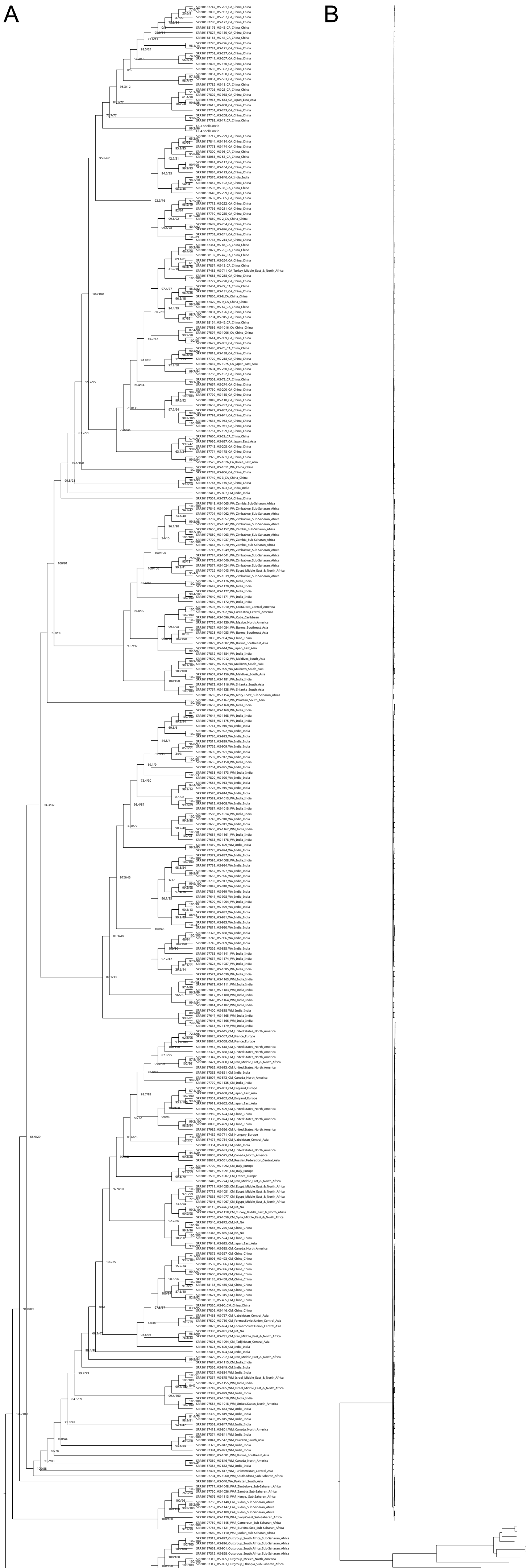


**Figure S7: Nuclear phylogenetic tree. a)** cladogram. Node labels give support values from SH-aLRT/UFBoot. **b)** phylogram with branch lengths in expected substitutions per site. Scale bar gives 1 expected substitutions per site.

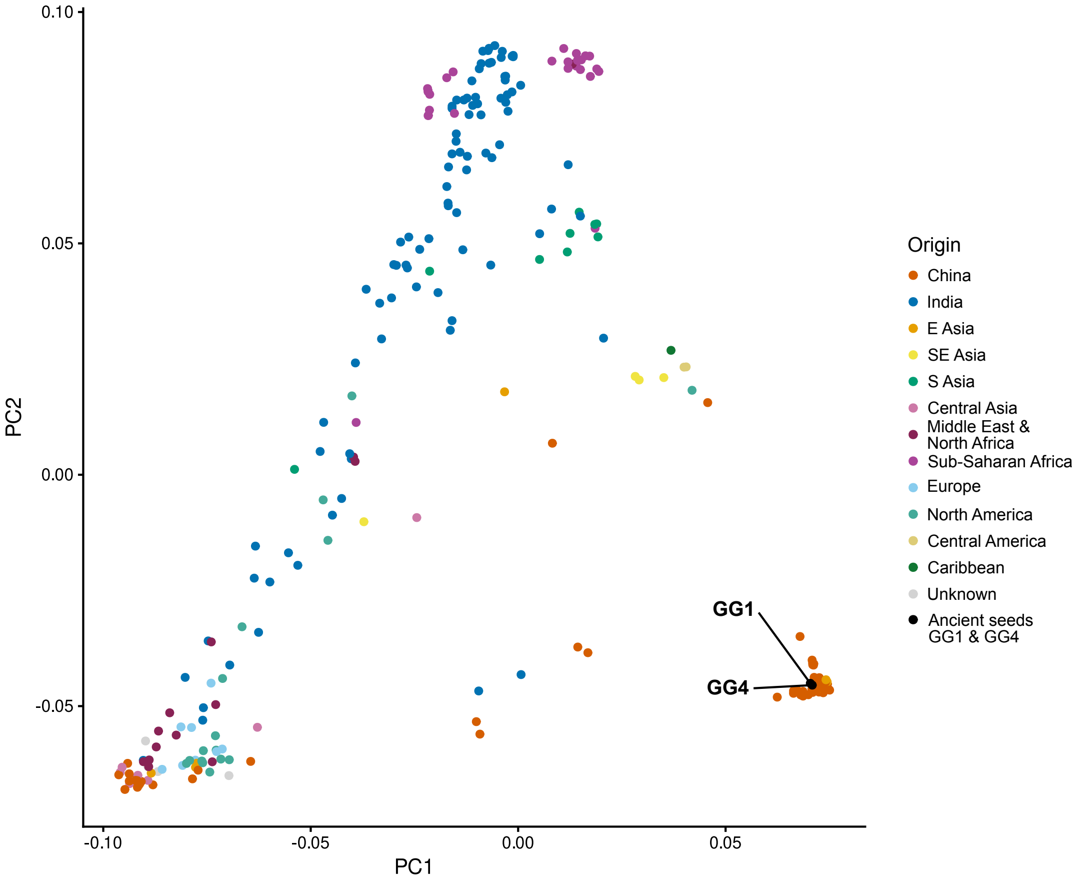


**Figure S8: PCA of nuclear genotype likelihoods from all positions.** Points are coloured by geographic origin of the accession and the two ancient seed genomes are labelled. Note that this run failed to converge in 1,000 iterations of PCANGSD.

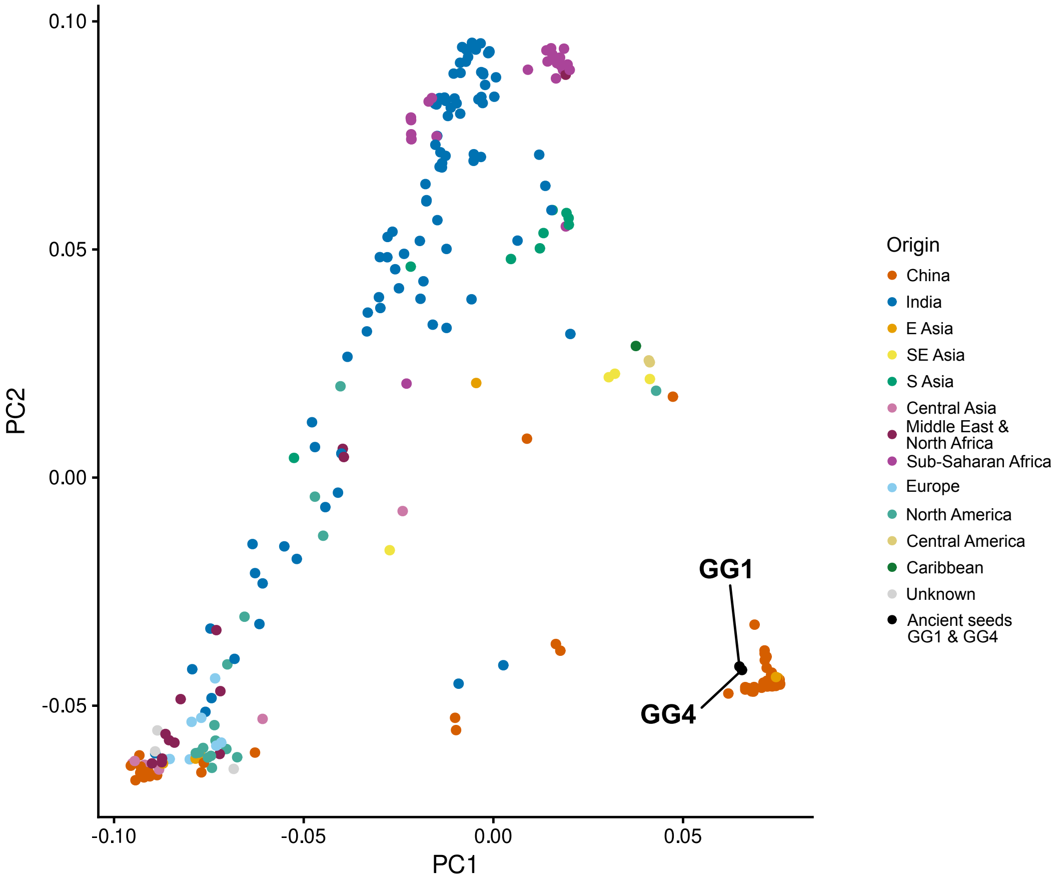


**Figure S9: EMU PCA of pseudohaploid genotype calls from all positions.** Points are coloured by geographic origin of the accession and the two ancient seed genomes are labelled.

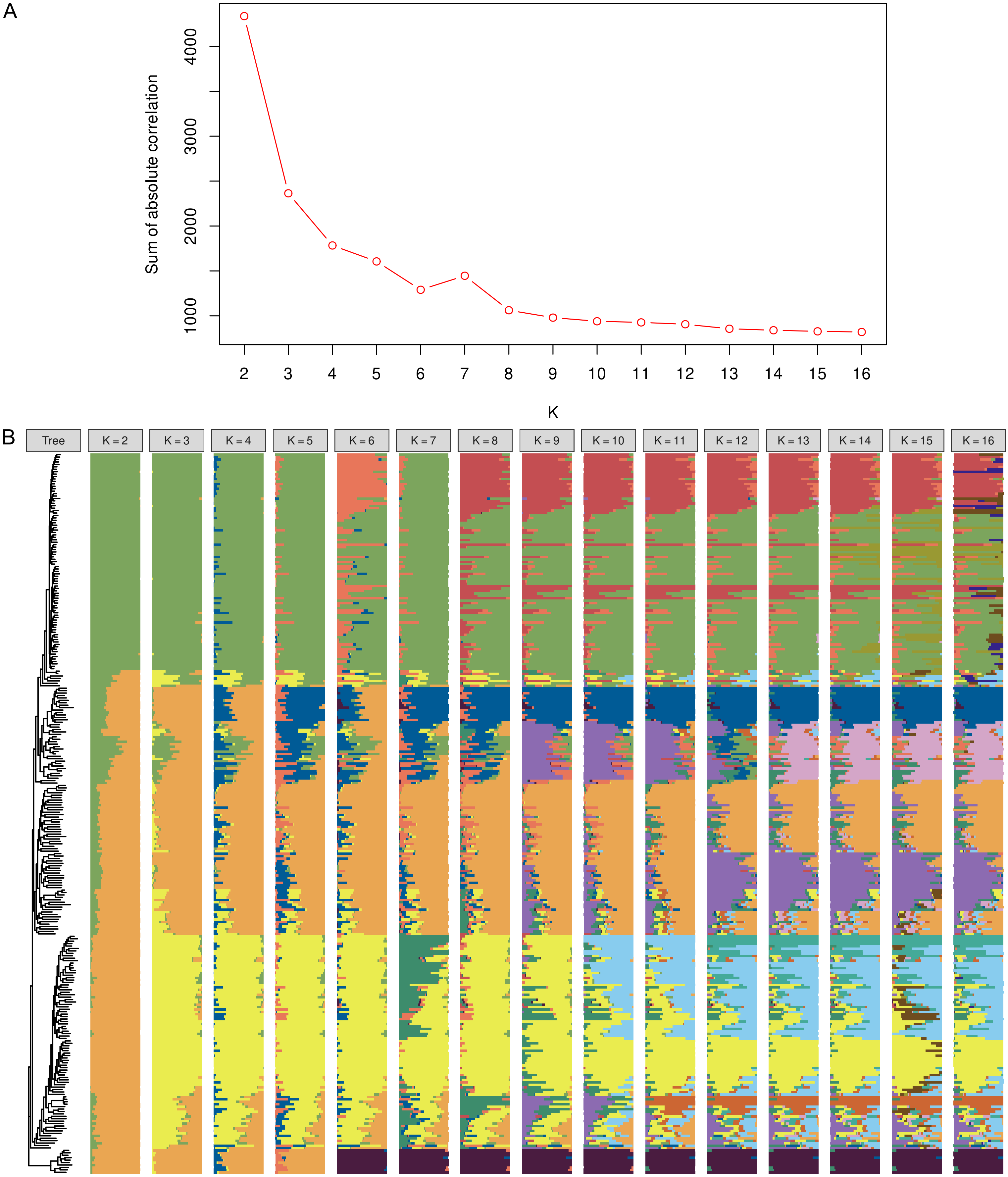


**Figure S10: Population structure analysis with NGSAdmix and evalAdmix.** Analysis conducted using the dataset with no missing data. **a)** sum of the absolute value of correlations of residuals for different values of K from evalAdmix using the admixture proportions estimated by NGSAdmix. Lower values show a better fit of the admixture model. **b)** Admixture proportions for K=2-16.

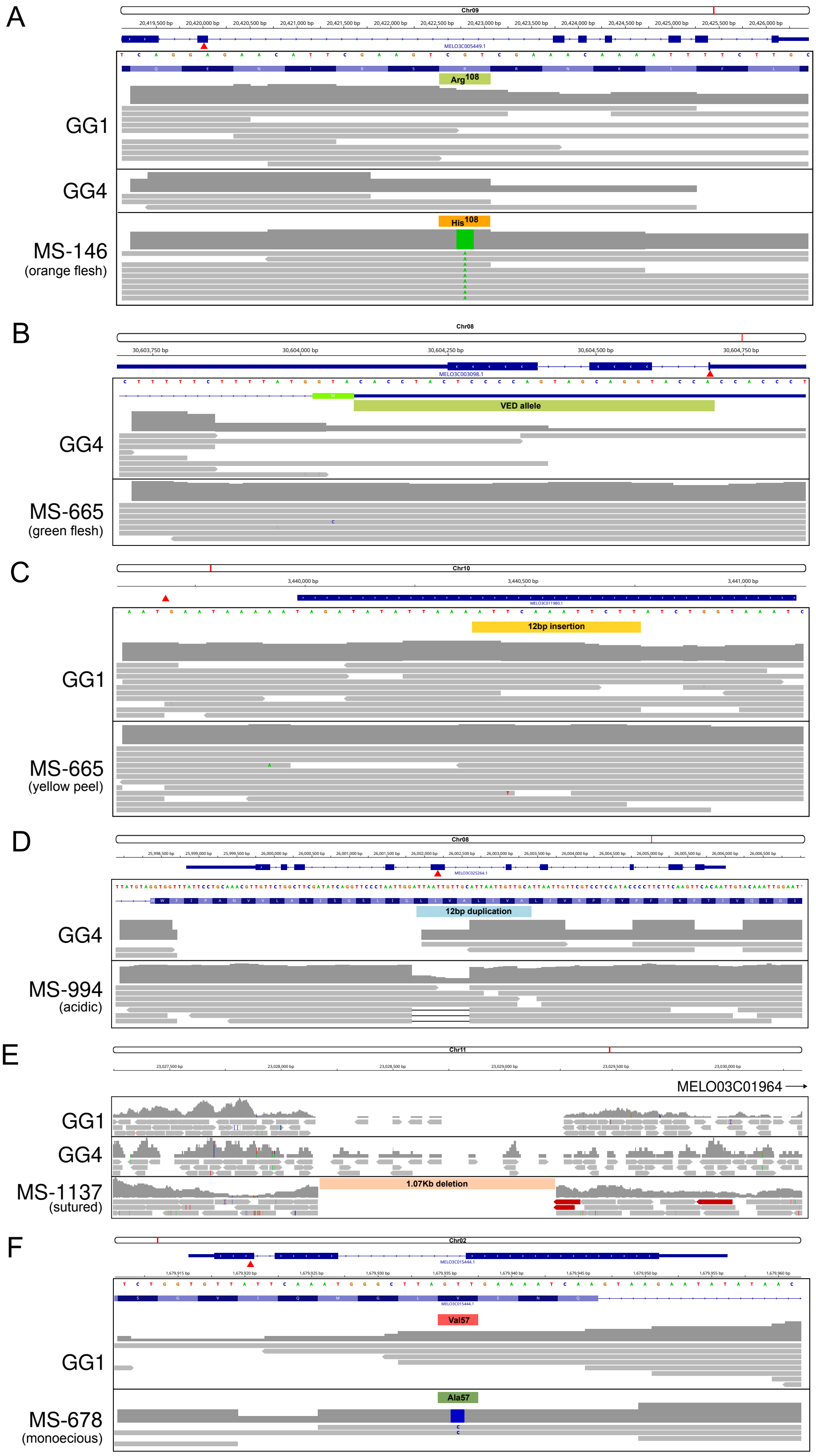


**Figure S11: Read pileups for key phenotypic loci. a)** CmOR **b)** CmRPGE1 **c)** CmKFB **d)** CmPH **e)** suture-associated deletion **f)** CmACS7
